# Gene expression profiles underlying aggressive behavior in the prefrontal cortex of cattle

**DOI:** 10.1101/2020.07.09.194704

**Authors:** Paulina G. Eusebi, Natalia Sevane, Manuel Pizarro, Marta Valero, Susana Dunner

## Abstract

Aggressive behavior is an ancient and conserved trait considered habitual and essential for most animals in order to eat, protect themselves from predators and also to compete for mating and defend their territories. Genetic factors have shown to play an important role in the development of aggression both in animals and humans, displaying moderate to high heritability estimates. However, although such types of conducts have been studied in different animal models, the molecular architecture of aggressiveness remains poorly understood. This study compared gene expression profiles of 16 prefrontal cortex (PFC) samples from aggressive and non-aggressive cattle breeds: Lidia, selected for agonistic responses, and Wagyu, specialized on meat production and selected for tameness. RNA sequencing was used to identify 918 up and 278 down-regulated differentiated expressed genes (DEG). The functional interpretation of the up-regulated genes in the aggressive cohort revealed enrichment of pathways such as the Alzheimer disease-presenilin, integrins or the ERK/MAPK signaling cascade, all implicated in the development of abnormal aggressive behaviors and neurophysiological disorders. Moreover, gonadotropins, leading to testosterone release, are also up-regulated as natural mechanisms enhancing aggression. Concomitantly, heterotrimeric G-protein pathways, associated with low reactivity mental states, and the *GAD2* gene, a repressor of agonistic reactions at PFC, are down-regulated, guaranteeing the development of the adequate responses required by the aggressive Lidia cattle. We also identified six upstream regulators, whose functional activity fits with the etiology of abnormal behavioral responses associated with aggression. These results provide valuable insights into the complex architecture that underlie naturally developed agonistic behaviors.

## Introduction

Aggressive behavior, an evolutionary well-conserved trait in animals, is part of the general conducts repertoire, as most animals need this skill in order to eat, protect themselves and their family against predators, compete for mating, as well as acquire resources and territory (Tremblay and Nagin, 2005). In contrast, aggressive behaviors in humans often refer to abnormal manifestations of aggressiveness such as violence, and are associated with a broad spectrum of neuropsychiatric disorders such as dementia, manic depression, bipolar disorder, or schizophrenia (Haller *et al*. 2006, Manchia *et al*. 2017). Research studies have shown that the expression of aggressive behavior depends on the interaction between environmental and genetic factors, with a genetic additive component ranging around 50% in humans (Tuvblad and Baker, 2011).

A large number of preclinical studies using different animal species as models has been encouraged on the reasoning that molecular correlates of animal aggressive behavior resemble biological mechanisms in human pathological aggression (Blanchard and Blanchard, 2003). Several attempts to mold abnormal forms of aggressiveness using mainly murine models, and to a lesser extent dogs and semi-domesticated species such as the silver fox, have been performed to display a contrast between docile or tame behaviors and escalated levels of aggressiveness (de Boer *et al*. 2003). However, relating these mechanisms to the human condition is not simple, since aggressive behaviors are very diverse. In animals, aggressive responses consist of a combination of fight, chase, bite and ram, whereas aggression in humans involves both verbal and physical forms. Despite this, it is possible to look for similarities between species in the components of aggression to better understand its etiology and to further improve its diagnosis, prognosis and intervention strategies, which currently lack in effectiveness (McGuire, 2008). In the bovine species, the Lidia breed may constitute a useful tool for studying the genomic makeup of aggressive behavior. Lidia bovines belong to a primitive population, selected for centuries to develop agonistic-aggressive responses by means of a series of traits registered by the breeders on a categorical scale that classifies their aggression and fighting capacity, reporting moderate to high heritability estimates (0.20 - 0.36) (Silva, Gonzalo & Cañón, 2006; Menéndez-Buxadera *et al*. 2017). A recent study has identified significant genomic regions containing genes associated with aggressive behavior in the Lidia breed (Eusebi *et al*. 2018), including a polymorphism in the promoter of the monoamine oxidase A (*MAOA*) gene, an important locus widely associated to pathological forms of aggression which, in humans, derives in a broad spectrum of psychiatric conditions, such as manic and bipolar disorders and schizophrenia, among others (Craig and Halton, 2009, Eusebi *et al*. 2019). However, no studies on gene expression differences for behavioral features have been conducted so far in cattle.

The genetic expression of behavior takes part in the brain, where the frontal cortical region, in particular the prefrontal cortex (PFC), has shown to play a crucial role in the regulation of aggressive behavior (Miczek *et al*., 2007; Siever, 2008). The PFC has been studied on different species, e.g. it has been suggested that PFC lesions result in impulsive and antisocial behaviors in humans (Brower *et al*. 2001) and offensive aggression in rodents (Craig and Halton, 2009). Moreover, a catalogue of gene-specific sequence variants was detected as differentially expressed between an aggressive-selected strain of silver fox when compared to its tamed counterpart (Kukekova *et al*. 2011). Similar results are reported in RNA-seq profiles of different dog breeds (Våge *et al*. 2010).

Thus, the goal of our study is to seek for genes that are differentially expressed in the PFC of aggressive and non-aggressive bovines using as models the Lidia and the Wagyu breeds for each cohort respectively. The two breeds differ significantly in their agonistic responses, the Lidia breed known as one of the most aggressive bovine breeds, whereas Wagyu bovines are calm and docile animals, selected and bred by farmers with the aim of easing their handling (Takanishi *et al*. 2015). This makes our studied populations highly suitable for investigating the biological underpinnings that may contribute to control naturally developed aggressive behavior in animals.

## Methods

No special permits were required to conduct the research. All animals were sacrificed for reasons other than their participation in this study. The access to the brains of the animals was allowed in the cutting room, after the “*corrida*” in the case of Lidia animals and after slaughter with the Wagyu breed animals, following standard procedures approved by the Spanish legislation applied to abattoirs and “*corrida*” festivities (Reglamento General de Mataderos/BOE-A-1997-3081).

### Animal and tissue processing

Post mortem PFC tissue samples were retrieved from 16 non-castrated male bovines aged 3 to 4 years, 8 belonging to the Lidia breed and 8 from the Wagyu breed, for the aggressive and non-aggressive cohorts respectively. The 8 Lidia bovines belong to two different lineages or *encastes*, Santa Coloma (N=4) and Domecq (N=4), as defined by the Boletín Oficial del Estado (2001). For approximately half an hour prior to its sacrifice (the time of the traditional “*corrida*” festivity), these bovines were incited to develop agonistic-aggressive behaviors by a series of responses that define their aggression and fighting capacity, as described by Domecq (2009). Non-aggressive Wagyu cattle samples were taken to the slaughterhouse where animals were transported a few hours before their sacrifice. The bovines were handled in-group, avoiding social encounters among them and agonistic reactions. The PFC tissue samples from both cohorts were harvested less than an hour post-mortem and immediately immersed in RNA-later™ (Thermo Fisher Scientific, Madrid, Spain), followed by 24 hours’ storage at 5°C and long-term conservation at -80°C.

### RNA extraction, sequencing and bioinformatics analyses

Total RNA was extracted from postmortem PFC tissue using the RNeasy Lipid Tissue Mini Kit (QIAGEN, Spain) according to the manufacturer’s instructions. Tissuelyser (QIAGEN, Spain) was used to homogenize samples. RNA quantification and purity were assessed with a Nanodrop ND-1000 spectophotometer (Thermo Fisher Scientific, Madrid, Spain) and RNA integrity number (RIN) was determined using the Bioanalyzer-2100 equipment (Agilent Technologies, Santa Clara, CA, USA). To guarantee its preservation, RNA samples were treated with RNAstable (Sigma-Aldrich, Madrid, Spain), and shipped at ambient temperature to the sequencing laboratory (DNA-link Inc. Seoul, Korea) to perform high throughput sequencing using a Novaseq 6000 sequencer (Illumina, San Diego, CA, USA). For quality check, the OD 260/280 ratio was determined to be between 1.87 and 2.0. Library preparation for Illumina sequencing was done using the Illumina Truseq Stranded mRNA Preparation kit (Illumina, San Diego, CA, USA). Sequencing was performed as 100 base paired-end mode, followed by automatic quality filtering following Illumina specifications. All these process where performed according to the manufacterer’s instructions. Individual reads were de-multiplexed using the CASAVA pipeline (Illumina v1.8.2), obtaining the FASTQ files used for downstream bioinformatics analysis. Illumina reads generated from all samples have been deposited in the NCBI GEO / bioproject browser database (Accession Number: GSE148938).

Read quality of the sixteen RNA-seq datasets was checked and trimmed using PRINSEQ v. 0.20.4 (Schmieder and Edwards, 2011). Trimmed reads were then mapped to the bovine reference genome (Bos taurus ARS.UCD 1.2) with STAR v.2.7.3a (Dobin et al. 2013), using default parameters for pair-end reads and including the Ensembl Bos taurus ARS-UCD 1.2 reference annotation. The SAM files generated by STAR, which contains the count of reads per base aligned to each location across the length of the genome, were converted into a binary alignment/map (BAM) format and sorted using SAMTools v.0.1.18 (Li et al. 2010). The aligned RNA-seq reads were assembled into transcripts and their abundance in fragments per kilobase of exon per million fragments mapped (FKPM) was determined with Cufflinks v.2.2.1 (Trapnell et al. 2010). The assembled transcripts of all samples were merged using the Cufflinks’ tool Cuffmerge. Analysis of differential gene expression across aggressive and non-aggressive groups was performed using Cuffdiff, included also in the Cufflinks package. A Benjamini-Hochberg False Discovery Rate (FDR), which defines the significance of the Cuffdiff output, was set as threshold for statistical significant values of the Differentially Expressed Genes (DEG). The R software application CummeRbund v.2.28.0 (Goff et al. 2012) was used to visualize the results of the RNA-seq analysis.

### Cross-species comparative analysis (CSCA)

Because no other differential expression analysis using cattle as animal model for aggressive behaviors has been conducted before, we performed a comparison among our DEG and a cross-species compendium of genes associated with aggressiveness previously identified in different studies in humans, rodents, foxes, dogs and cattle as proposed by Zhang-James *et al*. (2019). The gene compendium is a list based on four main categories of genetic evidence: i) two sets of genes identified in different genome-wide association studies (GWAS) in humans, one for adults and the other for children (Fernández del Castillo *et al*. 2016); ii) one set of genes showing selection signatures in Lidia cattle (Eusebi *et al*. 2018; Eusebi *et al*. 2019); iii) four sets of genes differentially expressed in rodents (Malki et al. 2014; Clinton *et al*. 2016) and one in silver foxes (Kukekova *et al*. 2011; Kukekova *et al*. 2018); and iv) three sets of genes with causal evidence from the Online Mendelian Inheritance in Man (OMIM) database, a knockout (KO) mice report and causal evidence in dogs retrieved from the Online Mendelian Inheritance in Animals (OMIA) database (Våge *et al*. 2010; Veroude *et al*. 2016; Zhang-James *et al*. 2019). To homogenize the compendium gene-list with our DEG, gene official names from cattle were converted to its human orthologues using biomaRt (Durinck *et al*. 2005). In order to establish a ranking according to the total occurrence of each gene in the different sets we assigned a weight (weighted ranking, WR) to each of our DEG in common with the compendium gene list applying the same conditions proposed by Zhang-James *et al*. (2019). The compendium gene-list and details of the different studies are described in Supplementary table 1.

### Gene ontology and KEGG pathway enrichment analyses

To examine the relationships between differences in PFC gene expression among groups and its biological function, we first separated the results of DEG in two independent gene-lists according to their Log_2_ Fold Change (FC): up-regulated for those transcripts displaying a Log_2_FC ≥ 0.1; and down-regulated for those with a Log_2_FC ≤ 0.1. The Panther database v.12.0 (http://www.pantherdb.org/) was then used to determine processes and pathways of major biological significance through the Over Representation test based on the Gene Ontology (GO) annotation function. Panther applies different algorithms using the uploaded reference lists as seeds and known interactions from the database and edges, to generate content specific pathways. We used the Fisher’s exact test for annotation and the FDR for multiple testing corrections, both for the up and down regulated DEG with P-values ≤ 0.05, to infer their pathway enrichment scores.

### Biological role of the genes in common with the CSCA: interactions and upstream regulators

The Ingenuity Pathway Analysis (IPA) (QIAGEN, www.qiagen.com/ingenuity) software was used to identify GOs, pathways and regulatory networks to which our DEG in common with the compendium gene-list belong to, as well as upstream regulators. IPA transforms a set of genes into a number of relevant networks based on comprehensive records maintained in the Ingenuity Pathways Knowledge Base. The Networks are presented as graphics depicting the biological relationships between genes and gene products. The analysis of upstream regulators considers all the possible transcription factors and upstream regulation, as well as their predicted effects on gene expression contained in the Base repository. Therefore, IPA allows to analyze if the patterns of expression observed in the DEG can be explained by the activation or inhibition of any of these regulators through an estimation of a z-score, which is a statistical measure of the match between the expected relationship direction among the regulator and its targets, and the observed gene expression (Krämer *et al*. 2014).

## Results

### Sequencing and read assembly

The RNA-sequencing of the sixteen PFC samples generated an average of 78.3 million paired-end reads per sample. The mean mapping proportions obtained with the STAR software was 91.8%, similar among different samples (from 88.07 to 94.91%) (Supplementary Table 2). The mapped reads were processed with Cufflinks toolkits for differential expression analysis, revealing a total of 16,384 DEG between the aggressive and non-aggressive groups; of those genes, 1196 were statistically significant, producing 10,640 isoforms (8.86 transcripts per gene) (Table 1, Figure 1A and Supplementary Figure 1). Gene expression differences of the up-regulated DEG (log_2_FC ≥ 0.1) were higher, involving 918 genes, than the down-regulated 278 DEG (log_2_FC ≤ 0.1) (Figure 1B and C). For the complete list of up and down-regulated DEG see Supplementary Table 3.

**Table 1.** Summary statistics of differentially expressed features.

| Classification | Transcripts |
| --- | --- |
| Significant DEG | 1,196 |
| Up-regulated DEG | 918 |
| Down-regulated DEG | 278 |
| Differentially expressed isoforms | 10,640 |

**Figure 1.**
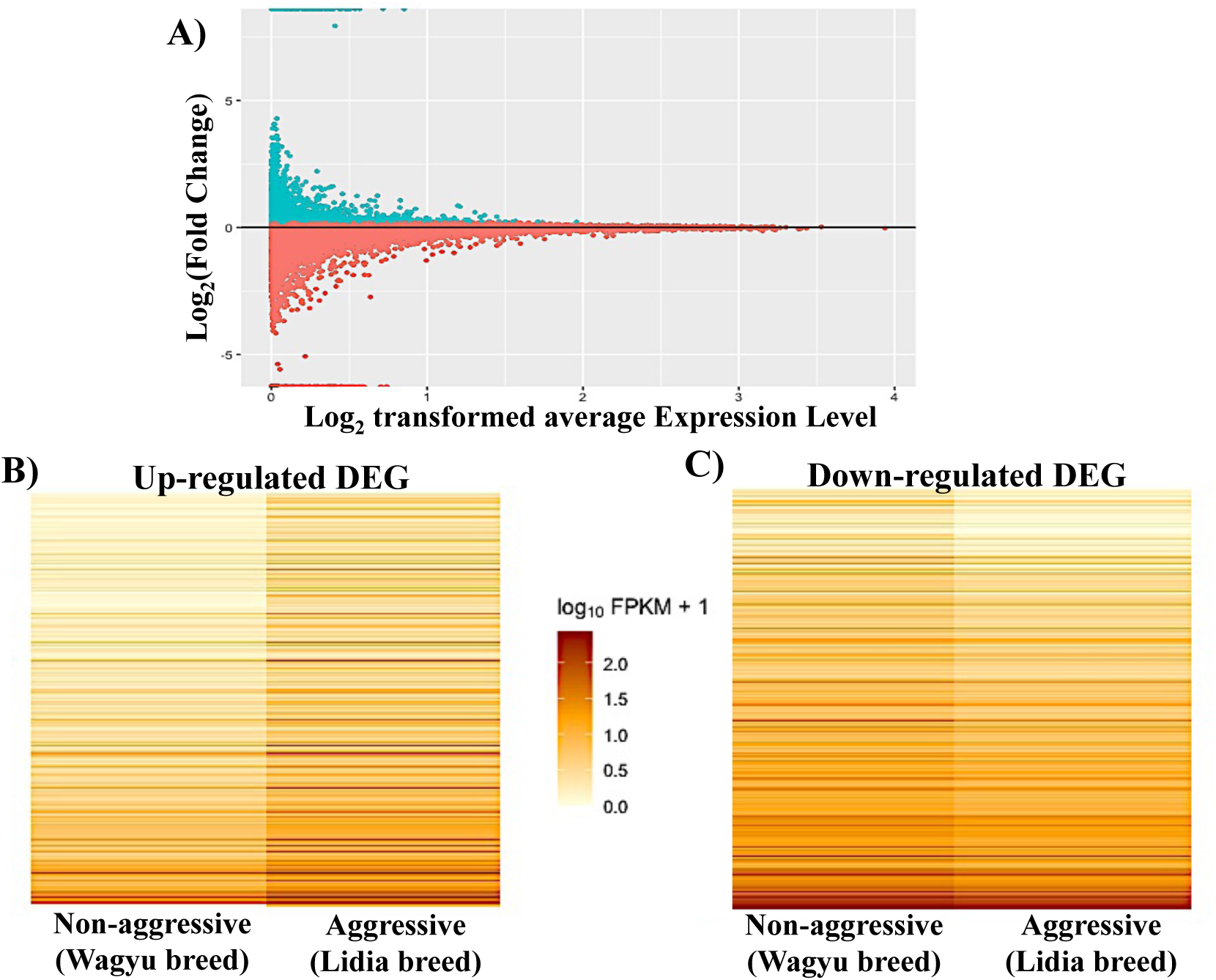
**A) MA-plot showing the distribution of differentially expressed genes (DEG).** The Y-axis shows the log_2_ (Fold Change) of expression between aggressive and non-aggressive groups, and the X-axis correspond to the log_2_ transformed average expression level for each gene across samples. Log_2_FC ≥ 0.1 and Log_2_FC ≤ 0.1 genes are represented by green and red dots, respectively. **B) Heatmap of up-regulated DEG** in the aggressive group. **C) Heatmap of down-regulated DEG** in the aggressive group.

### Genes in common with the cross-species comparative analysis (CSCA)

The up and down-regulated DEG were compared with the compendium genes-list associated with aggressive behavior (Supplementary Table 1). This comparison yielded 31 genes, 17 up and 14 down-regulated in the aggressive group of Lidia individuals (Table 2). The expression bar plot in Figure 2 shows the common subset of DEGs in relation to their standard deviation FPKM values and their WR. Most of the overlapping genes from the up-regulated DEG coincide with GWAS on aggressive behaviors in humans. Instead, the down-regulated DEG greater coincidences were detected with differential expression analyses and KO studies in mice.

**Figure 2.**
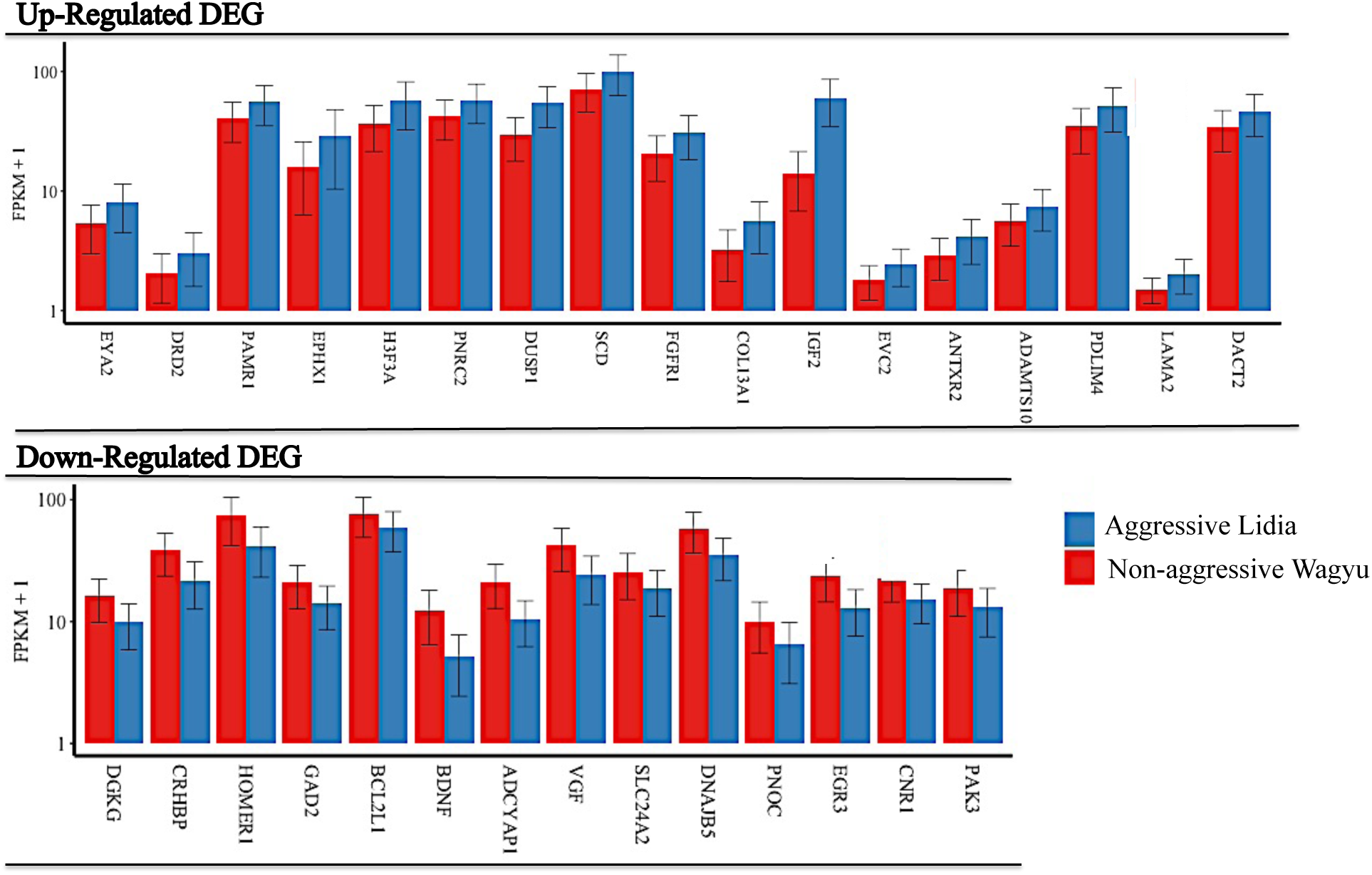
Bar chart of up and down regulated DEG in common with the cross-species comparative analysis (CSCA). Gene abundance is represented in fragments per kilobase of exon per million fragments mapped (FPKM) of the aggressive (Lidia breed) and non-aggressive (Wagyu breed) cohorts.

**Table 2.** Up and down regulated DEG in common with the cross-species comparative analysis (CSCA).

| UP-REGULATED |  |  |
| --- | --- | --- |
| Gene symbol | Gene name | Weighted Ranking (WR) |
| *LAMA2* | ADAM metalloproteinase with thrombospondin type 1 motif 1 | 2 |
| *DRD2* | Dopamine receptor 2 | 2 |
| *DUSP1* | Dual specificity phosphatase 1 | 1.5 |
| *FGFR1* | Fibroblast growth factor receptor 1 | 1 |
| *PDLIM4* | PDZ and LIM domain 4 | 1 |
| *PNRC2* | Proline rich nuclear receptor co-activator 2 | 1 |
| *SSD* | Stearoyl-CoA desaturase | 1 |
| *H3F3A* | H3.3 Histone A | 1 |
| *PAMR1* | Peptidase domain containing associated with muscle regeneration 1 | 1 |
| *ADAMTS1* | ADAM metalloproteinase with thrombospondin type 1 motif 1 | 1 |
| *ANTRX2* | ANTRX cell adhesion molecule 2 | 1 |
| *COL13A1* | Collagen type XIII alpha chain 1 | 1 |
| *DACT2* | Dishevelled binding antagonist of beta catenin 2 | 1 |
| *EPHX1* | Epoxide hydrolase 1 | 1 |
| *EVC2* | EvC ciliary complex subunit 2 | 1 |
| *EYA2* | EYA Transcriptional co-activator and phosphatase 2 | 1 |
| *IGF2* | Insulin like growth factor 2 | 1 |

| DOWN-REGULATED |  |  |
| --- | --- | --- |
| Gene symbol | Gene name | Weighted Ranking (WR) |
| *GAD2* | Glutamate decarboxylase 2 | 2.5 |
| *PNOC* | Propionociceptin | 2.5 |
| *DNAJB5* | DnaJ Heat shock protein family (Hsp40) member B5 | 2 |
| *ADCYAP1* | Adenylate cyclase activating polypeptide 1 | 2 |
| *BDNF* | Brain derived neurotrophic factor | 2 |
| *CRHBP* | Corticotropin releasing hormone binding protein | 2 |
| *EGR3* | Early growth response 3 | 2 |
| *HOMER1* | Homer scaffold protein 1 | 2 |
| *PAK3* | P21 (RAC1) activated kinase 3 | 2 |
| *BCL2L1* | BCL2 like 1 | 1.5 |
| *DGKG* | Diacylglycerol kinase gamma | 1.5 |
| *SLC24A2* | Solute carrier family 24 member 2 | 1 |
| *VGF* | VGF nerve growth factor inducible | 1 |

### Functional annotation and biological pathway analysis

A GO analysis of the pathways and biological processes identified in the data set lists containing significant up and down-regulated transcripts was carried out. Among the 918 up-regulated DEGs in aggressive Lidia samples, Panther Over Representation test included 851 uniquely mapped IDs, displaying significant association with 881 GO biological processes (FDR ≤ 0.05), most of them related to heart morphogenesis and heart development, cellular adhesion, migration and differentiation, skeletal and smooth muscle development, central nervous system (SNC) development and function, and immune response (Supplementary Table 4). The Panther Pathway enrichment analysis retrieved five significant pathways: blood coagulation, integrin signaling, Alzheimer disease-presenilin, angiogenesis and gonadotropin-releasing hormone receptor pathways (Table 3A). Within the down-regulated DEGs in the aggressive cohort, the GO biological processes included 260 genes as uniquely mapped IDs implicated in 243 processes (FDR ≤ 0.05), the higher significant values, being dendritic cell cytokine production, trans-synaptic signaling by endocannabioid, trans-synaptic signaling by lipid, negative regulation of renin secretion into blood stream and melanocyte adhesion, all with 84.4 fold enrichment and two genes associated with each process (Supplementary table 5). The Panther enrichment pathway analysis retrieved two significant down-regulated pathways in the aggressive Lidia breed, both involved in two different types of Heterotrimeric G-protein signaling (Table 3B).

**Table 3.** A) Panther enriched pathways of the up-regulated differentially expressed genes (DEG) in the aggressive Lidia breed.

| Pathway | DEG | Fold Enrichment | P-value | FDR |
| --- | --- | --- | --- | --- |
| Blood coagulation | 10 | 4.32 | 2.51E-04 | 8.33E-03 |
| Integrin signaling pathway | 29 | 3.85 | 5.47E-09 | 4.54E-07 |
| Alzheimer disease-presenilin pathway | 18 | 3.67 | 7.66E-06 | 4.24E-04 |
| Angiogenesis | 21 | 3.15 | 1.18E-05 | 4.92E-04 |
| Gonadotropin-releasing hormone receptor pathway | 23 | 2.26 | 7.08E-04 | 1.96E-02 |

**Table 3.** B) Panther Enriched pathways of the down-regulated DEG in the aggressive Lidia breed.

| Pathway | DEG | Fold Enrichment | P-value | FDR |
| --- | --- | --- | --- | --- |
| Heterotrimeric G-protein signaling pathway-Gq alpha and Go alpha mediated pathway | 8 | 5.32 | 1.96E-04 | 1.63E-02 |
| Heterotrimeric G-protein signaling pathway-Gi alpha and Gs alpha mediated pathway | 9 | 4.34 | 3.33E-04 | 1.84E-02 |

### Signaling networks and upstream regulators enrichment analysis

We used the IPA software to identify pathways to which DEGs in common with the CSCA belong to, as well as to explore the existence of signaling networks connecting the DEGs. Sixty-eight categories with P-values ≤ 0.05 were associated with different diseases or function annotations significantly enriched in the 31 common genes from the CSCA (Supplementary Table 6). This information should be interpreted cautiously because most pathways were represented by a small number of genes (from 1 to 5).

The top-scoring regulatory network identified predicted interactions among 35 genes, including three up and two down-regulated DEGs in the aggressive cohort, and is related with functions such as behavior, cellular movements and embryonic development (Figure 3). Amongst the diseases and functional annotations, it is worth to mention aggression (P-value = 1.37E-05), learning behavior (2.24E-04), memory (2.48E-04), cognition (3.46E.04) brain damage (9.41E-04), progressive encephalopathy (1.08E-03), schizophrenia (1.22E-03), forgetting (1.91E-03), and processes involving nervous system development and formation (Supplementary Table 6).

**Figure 3.**
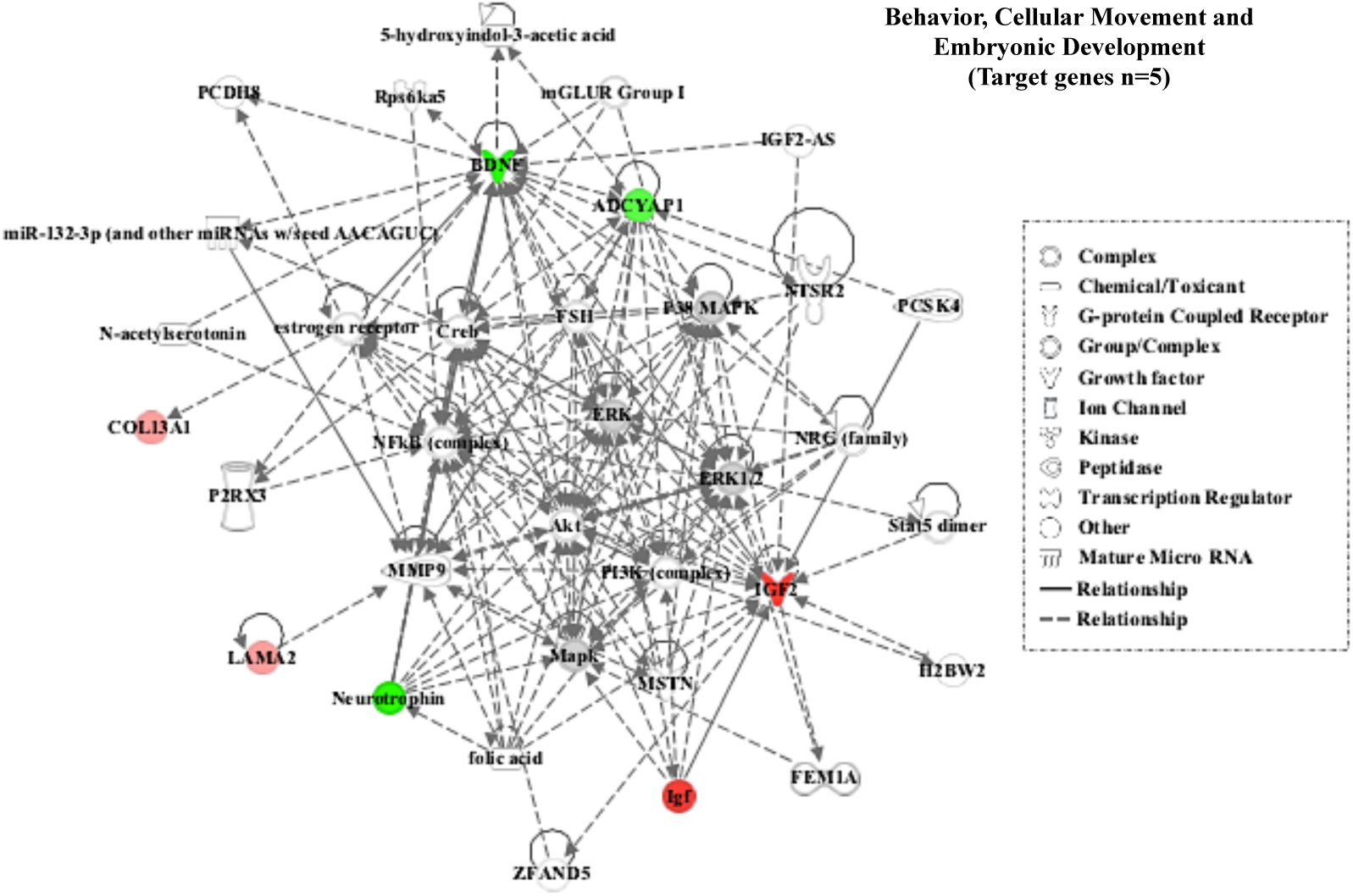
Top-scoring regulatory network identified with the IPA software corresponding to behavior, cellular movements and embryonic development functions. Up and down-regulated differentially expressed genes (DEG) in the aggressive Lidia breed are displayed with red and green nodes, respectively. Genes are represented as nodes, and the molecular relationship between nodes is represented either as straight lines for direct interactions, or dotted lines for indirect interactions.

Finally, the upstream analysis tool of the IPA package was used to identify the potential upstream regulators that may explain the differential patterns of expression between the up and down regulated DEGs in common with the CSCA in the aggressive cohort. By doing so, six main upstream regulators were identified: Insulin-Like Growth factor 2-Antisense RNA (IGF2-AS), Neurotrophic Receptor Tyrosine Kinase 1 (*NTRK1*), Amyloid Beta Precursor Protein (*APP*), RAD21 Cohesin complex component (*RAD21*), Zinc finger BED-Type Containing 6 (*ZBDE6*), and Hedgehog (Hh) (Figure 4). All these genes, RNAs and proteins appear to be involved in a heterogeneous array of biological functions related to behavior development and cell-to-cell signaling interactions.

**Figure 4.**
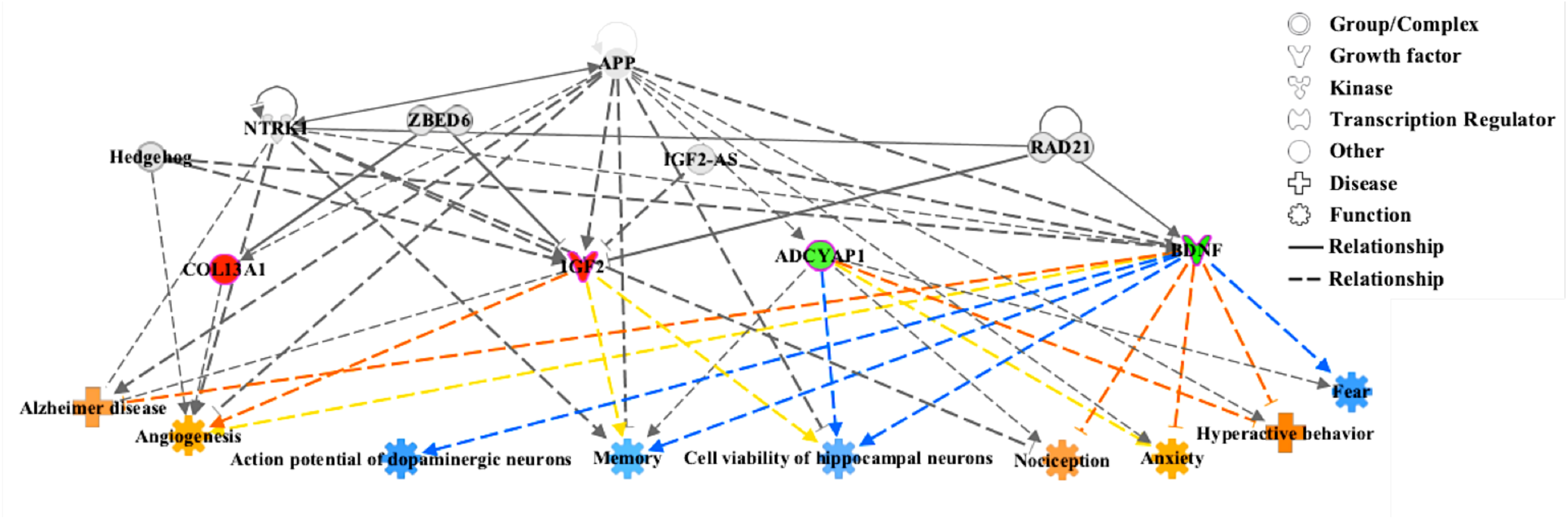
Major upstream regulators of the network of the differentially expressed genes (DEG) in the aggressive Lidia group. There are six up-stream regulators predicted to be activated (gray color). In red (up-regulated) and green (down-regulated) we can see the genes whose expression changes in response to the activation of the upstream regulators. The shapes of the nodes represent the functional class of each gene or gene product, as defined in the legend. The straight and dashed lines represent direct and indirect interactions, orange leads to activation, blue leads to inactivation, gray is an effect not predicted and yellow are inconsistent findings with the state of the downstream molecule.

## Discussion

Understanding the complexity of the mechanisms behind the development of aggressive behaviors in humans and animals is still a challenge, although several molecular studies using different animal models apart from humans addressed this goal in the last years (Clinton *et al*. 2016; Eusebi *et al*. 2018; Eusebi *et al*. 2019; Fernández del Castillo *et al*. 2016, Kukekova *et al*. 2011; Kukekova *et al*. 2018; Malki *et al*. 2014; Våge *et al*. 2010; Veroude *et al*. 2016; Zhang-James *et al*. 2019). The present study represents the first description of transcriptional mechanisms affecting aggressive behavior in cattle.

The number of DEG identified in the bovine PFC (16,384) is similar to that identified in mice (15,423) (Han *et al*. 2009) and silver fox (14,000) (Kukekova *et al*. 2018), also in PFC. After correcting for Log_2_ Fold Change (FC), 918 up and 278 down regulated genes displayed a wide array of functional pathways. Within the up-regulated enriched pathways in the aggressive cohort, we found biological functions related with processes such as cellular, muscular and SNC development and function, heart formation and development, and immune responses (Supplementary Table 4). Similar results were obtained by Kukekova *et al*. (2011); they compared the PFC expression between aggressive and docile strains of silver fox and also observed an enrichment of pathways related to cellular movement, growth and proliferation, hematological system development and antigen presentation.

Among the top enriched up-regulated pathways in the aggressive Lidia group, the integrin and the Alzheimer disease-presenilin signaling pathways were well-known gene routes in the development of abnormal aggressive behaviors (Wu *et al*. 2012) (Table 2A). At nervous system, integrins are essential molecules to brain’s neuroplasticity, i.e. the ability to adapt to internal and external stimuli by reorganizing its structure, function and connections (Cramer *et al*. 2011). Integrins increased expression contributes to imbalanced synaptic function in specific pathological conditions, such as Alzheimer disease and schizophrenia, both often accompanied by episodic aggression and violence (Wu *et al*. 2012). It has been detected that presenilin aberrant expression also plays an important role in Alzheimer, with behaviors as agitation and aggression being frequently occurring symptoms for individuals with this disease (Siever, 2008; Ballard *et al*. 2009).

Also, the overexpression of genes belonging to the gonadotropin-realizing hormone (GnRH) receptor pathway may have a strong impact on the biological mechanisms leading to aggression. Wise *et al*. (2000) observed that, in boars, increased serum concentrations of GnRH results in higher levels of testosterone. Testosterone is a sex hormone that has been implicated in the modulation of PFC; when increased, it may affect the fear-processing circuitry, which has been associated with reactive and abnormal aggression responses (Radke *et al*. 2015; Bakker-Huvenaars *et al*. 2018).

Curiously, the up-regulated pathway showing the highest over-representation in the group of animals displaying agonistic behavior (4.32 fold enrichment), includes genes associated with blood coagulation. The links between the blood coagulation system and behavior are being increasingly recognized. For example, Yang *et al*. (2016) observed a strong association of genes belonging to the blood coagulation pathway in human psychiatric disorders, such as major depression and suicidal behavior. The interrelation of hemostasis and angiogenesis, with the regulation of angiogenesis during vessel repair mediated by proteins secreted by platelets (Browder *et al*. 2000), may explain the concomitant up-regulation of the angiogenesis pathway found here.

Regarding the down-regulated DEG detected in the group of aggressive animals, we found the heterotrimeric G-protein pathways strongly suppressed (Table 2B). These routes are the main signaling pathways downstream receptor activation and, in this context, its functional status has been associated with major depression and bipolar disorders (González-Maeso and Meana, 2006). The fighting reaction that bulls elicit in a *corrida*, may temporarily antagonize the mechanisms implicated in low reactivity mental states, similar to those described in major depression disorder (González-Maeso and Meana, 2006).

To further disentangle the mechanisms activated by agonistic behaviors, we compared our dataset of DEG with those reported by Zhang-James *et al*. (2019) in humans and mice, Kukekova *et al*. (2011) in silver fox, Eusebi *et al*. (2018 and 2019) in cattle and Våge *et al*. (2010) in dogs. As shown in Figure 2, the level of concordance was quite low (only 31 genes in common were identified). Similarly, Zhang-James *et al*. (2019) reported a modest gene overlap between different categories of genetic evidence (human GWAS, genes with known causal evidence and rodent transcriptome genes). According to this author, the lack of overlap between studies suggests differences in the genetic etiology of aggression in different species and populations, and supports the complementarity of the gene sets detected.

All the 17 up-regulated genes associated with aggressiveness and shared with previous studies, are essential for neurodevelopment. The highest weight ranked the Laminin Subunit Alpha 2 (*LAMA2)* gene, which encodes an extracellular matrix protein and its mutation derives in a denervation atrophy of the muscle (Hall *et al*. 2007). The D2 dopamine receptor (*DRD2*) is one of the top ranked genes and has been widely studied on schizophrenia, for which SNPs located in the gene promoter affect its transcriptional activity (d’Souza *et al*. 2003). Among the 14 down-regulated genes in common with the CSCA, a notable finding concerns the Glutamate Decarboxylase 3 (*GAD2*) one of our highly ranked genes which is considered a “top-down” modulator of aggressive acts, playing a pivotal role in the control systems deployed by the PFC to moderate agonistic reactions (Jager *et al*. 2017). This gene, is a Gamma-aminobutyric acid type A (GABA)-synthesizing enzyme that has an inverse but linear relationship with aggression measures, meaning that low levels of GABBA are associated with high levels of aggression (Jager *et al*. 2017).

The analysis of the data with the IPA upstream enrichment tool retrieved one regulatory network related with behavior, cellular movements and embryonic development functions. In the network shown in Figure 3, two Mitogen Activated Protein Kinases (*MAPK*) and two Extracellular Signal Regulated Kinases (*ERK*) (outlined in grey color) occupy a central position. Similar results were obtained by Zhang-James *et al*. (2019), who identified also the ERK/MAPK signaling as mechanisms underlying aggression. Malki *et al*. (2014) performed a genome-wide transcriptome analysis of mouse models of aggression and also observed that the MAPK signaling pathway was differentially expressed between the aggressive and non-aggressive lines. The *MAPK/ERK* cascade is a key regulator of cell growth and proliferation, but most important, this signaling pathway activates the binding of different integrins at the cell surface to extracellular matrix proteins (Yee *et al*. 2008), linking its function with the up-regulated integrin pathway explained above. Finally, six upstream regulators were predicted to be major transcriptional regulators of the set of four DEG detected, *COL13A1, IGF2, ADCYAP1* and *BDNF* (Figure 4). The modulator effect of these molecules appears to increase the up-regulation of biological processes such as hyperactive behavior and anxiety, which are often associated with aggressiveness, and Alzheimer disease, a concordance feature with the above findings. We also found that the upstream regulators promote an increased nociception ability. Although it has never been implicated in aggression, it makes sense for Lidia cattle to display an enhancement of the capacity to respond to potentially damaging stimuli, and hence, display aggressive behaviors.

In conclusion, this is the first time a comparison of the differences in genomic expression between aggressive and non-aggressive selected cattle breeds at PFC has been performed, identifying 918 up and 278 down-regulated genes. We have also undertaken a cross-species comparison analysis to identify genes in common implicated in aggressiveness and investigate their regulatory networks. Our results include the up-regulation in the aggressive cohort of animals of pathways such as the Alzheimer disease-presenilin, integrins or the ERK/MAPK signaling cascade, all routes implicated in the development of abnormal aggressive behaviors and neurophysiological disorders, as well as normal mechanisms enhancing aggression such us the up-regulation of gonadotropins and, hence, testosterone, whose levels have been widely linked with agonistic reactions. On the contrary, heterotrimeric G-protein pathways, previously associated with low reactivity mental states like those involved in major depression, or the *GAD2* gene, with a pivotal role in the control systems deployed by the PFC to repress agonistic reactions, are both down-regulated, guaranteeing the development of the adequate combative responses needed during a “*corrida*” festivity. However, despite PFC is a key region for the modulation of aggressive behavior, it may not be representative of other brain regions reported also to play important roles in aggression, such as the hippocampus or the hypothalamus. Nevertheless, this constitutes the first important step towards the identification of the genes that impulse aggression in cattle and, by doing so, we are providing a novel species as model organism for disentangling the mechanisms underlying variability in aggressive behavior.

## Acknowledgements

The authors are grateful to the enterprise of Las Ventas, bullring “*Plaza 1*” and its veterinarian’s team for providing post-mortem Lidia breed brain tissue samples for this study. We also thank to *Nuestro Buey* (https://www.fincasantarosalia.com/)for providing the Wagyu breed post-mortem samples. Authors declare they do not have conflict of interests.

## Availability of data

Illumina reads generated from all samples have been deposited in the NCBI GEO / bioproject browser database (Accession Number: GSE148938).

## Supporting Information

**Figure Supplementary 1**. Isoforms (10,640) of the significant transcripts (1,1996). Values on the Y axis are differences in the squared coefficient of variation (CV^2^), a normalized measure of cross-replicate variability useful for evaluating the quality of the RNA-seq data. The X axis is the variability between replicate of fragments per kilobase of exon per million fragments mapped (FPKM) estimates.

## Notes

### Competing Interest Statement

The authors have declared no competing interest.

